# De novo purine metabolism is a metabolic vulnerability of cancers with low p16 expression

**DOI:** 10.1101/2023.07.15.549149

**Authors:** Naveen Kumar Tangudu, Raquel Buj, Hui Wang, Jiefei Wang, Aidan R. Cole, Apoorva Uboveja, Richard Fang, Amandine Amalric, Peter Sajjakulnukit, Maureen A. Lyons, Kristine Cooper, Nadine Hempel, Nathaniel W. Snyder, Costas A. Lyssiotis, Uma R. Chandran, Katherine M. Aird

## Abstract

p16 is a tumor suppressor encoded by the *CDKN2A* gene whose expression is lost in ∼50% of all human cancers. In its canonical role, p16 inhibits the G1-S phase cell cycle progression through suppression of cyclin dependent kinases. Interestingly, p16 also has roles in metabolic reprogramming, and we previously published that loss of p16 promotes nucleotide synthesis via the pentose phosphate pathway. Whether other nucleotide metabolic genes and pathways are affected by p16/*CDKN2A* loss and if these can be specifically targeted in p16/*CDKN2A*-low tumors has not been previously explored. Using CRISPR KO libraries in multiple isogenic human and mouse melanoma cell lines, we determined that many nucleotide metabolism genes are negatively enriched in p16/*CDKN2A* knockdown cells compared to controls. Indeed, many of the genes that are required for survival in the context of low p16/*CDKN2A* expression based on our CRISPR screens are upregulated in p16 knockdown melanoma cells and those with endogenously low *CDKN2A* expression. We determined that cells with low p16/*Cdkn2a* expression are sensitive to multiple inhibitors of *de novo* purine synthesis, including anti-folates. Tumors with p16 knockdown were more sensitive to the anti-folate methotrexate *in vivo* than control tumors. Together, our data provide evidence to reevaluate the utility of these drugs in patients with p16/*CDKN2A*-low tumors as loss of p16/*CDKN2A* may provide a therapeutic window for these agents.

## Introduction

p16, encoded by the *CDKN2A* locus, is a critical cell cycle regulator that inhibits CDK4/6 (Sherr, 2001). Given its importance in the cell cycle, p16 is a well-known tumor suppressor with ∼50% of all human tumors showing decreased expression or activity of p16 through a variety of mechanisms, including hypermethylation or deletion of the *CDKN2A* locus or mutations (Esteller et al., 2001; Inoue and Fry, 2018; Ruas et al., 1999). While the CDKN2A gene also encodes p14, cancer-associated mutations are more commonly found in p16 (Quelle et al., 1997), suggesting a more critical regulatory role of p16 in cancer. In melanoma, ∼30% of patients have a deep deletion of the *CDKN2A* locus and another 10-15% have *CDKN2A* mutations (Zhao et al., 2016). In spite of this, there are currently no FDA-approved therapies that are used specifically for these patients, although clinical trials of CDK4/6 inhibitors are ongoing in combination with both mutant BRAF inhibitors and immune checkpoint blockade (Garutti et al., 2021; Lelliott et al., 2022) (clinicaltrials.gov).

We and others have published that p16 has roles beyond cell cycle control, and some of these non-canonical roles are related to metabolism (Buj and Aird, 2019). We published that loss of p16 expression promotes nucleotide synthesis at least in part via increased translation of the pentose phosphate pathway (PPP) enzyme ribose-5-phosphate isomerase A (RPIA) (Buj et al., 2019). However, glucose tracing through the PPP indicated that additional pathways may contribute to the increase in steady state nucleotide levels observed in these cells. Moreover, there is currently no available RPIA inhibitor. Therefore, the goal of this study was to identify additional targetable nucleotide metabolism pathways that are specifically vulnerable to inhibition in p16-low cancers. We used melanoma as a model since ∼40% of melanomas have decreased p16, typically due to 9p21 chromosomal loss (Zhao et al.).

Nucleotide homeostasis is critical for cellular fidelity as nucleotides are not only used for energy and biomass, but also as signaling molecules and to cope with DNA damage (Aird and Zhang, 2015; Ali and Ben-Sahra, 2023; Shi et al., 2023; Turgeon et al., 2018). As such, multiple metabolic pathways feed into *de novo* nucleotide synthesis, and nucleotides can also be salvaged through catabolism or the microenvironment (Buj and Aird, 2018; Kohnken et al., 2015; Lane and Fan, 2015; Tong et al., 2009). Glucose via the PPP is used to synthesize phosphoribosyl pyrophosphate, the pentose sugar backbone of all nucleotides. One carbon metabolism is also critical, especially for *de novo* purine synthesis as purine carbons are derived from serine via the folate cycle (Yang and Vousden, 2016). Other metabolites that contribute to *de novo* synthesis include aspartate, glycine, and glutamine, thereby connecting a variety of metabolic pathways with nucleotide biosynthesis (Tong et al., 2009). It is well-known that many of these pathways are highly upregulated in cancer cells. Indeed, anti-metabolites, and specifically anti-folates, were some of the first used chemotherapies, and many are still used today. Whether these drugs could specifically benefit patients with p16-low tumors has not been investigated.

Using CRISPR KO libraries in both human and mouse isogenic p16 wildtype and shp16/sh*Cdkn2a* cells, in addition to mining the Dependency Map (DepMap), we identified 31 common nucleotide metabolism genes that are negatively enriched in p16/*CDKN2A*-low cells. Many of these genes are either transcriptionally or translationally upregulated in p16/*CDKN2A*-low cells. Therapeutic targeting of pathways that promote *de novo* purine synthesis, including anti-folates, demonstrated that p16/*Cdkn2a*-low cells are vulnerable to inhibition of these pathways *in vitro*. Treatment of mice bearing shp16 tumors with the anti-folate methotrexate decreased tumor burden that was not associated with toxicity. *In vitro*, treatment with anti-folates had a more robust long-term inhibitory effect compared to the CDK4/6 inhibitor palbociclib. Finally, mining of publicly-available data demonstrated gene alterations in a variety of enzymes related to nucleotide synthesis, although only those involved in purine synthesis and one carbon metabolism are associated with worse patient survival. Together, our data suggest p16/*CDKN2A* loss may create a therapeutic window to kill cancer cells with widely-used anti-folates with relatively little toxicity.

## Results

### CRISPR dropout screens identify multiple nucleotide metabolism vulnerabilities in shp16/sh*Cdkn2a* cells

We previously published that knockdown of p16 increases nucleotide synthesis via translation of the PPP enzyme RPIA (Buj et al., 2019). There are no available RPIA inhibitors; thus, we set out to identify additional nucleotide metabolic enzymes that are vulnerabilities of shp16 cells. We constructed a focused sgRNA library of 128 genes involved in nucleotide biosynthesis, salvage, and catabolism (**Table S1**) and performed a loss-of-function screen in SKMEL28 human melanoma cells with wildtype p16 and p16 knockdown (**Fig. 1A and S1A**) using a hairpin that we have previously characterized that can be rescued by p16 overexpression, suggesting a lack of off-target effects (Buj et al., 2019). Notably, this hairpin only targets p16 and not p14 (hairpin information in Materials and Methods). As expected, these cells proliferate faster than isogenic controls (**Fig. S1B**). Using MAGeCK analysis (Li et al., 2014), we observed negative selection for many genes involved in nucleotide homeostasis (**Fig. 1B and Table S2**). This included *RPIA*, demonstrating the validity of our screen (**Fig. S1C**). We next performed a secondary drop out screen in a mouse melanoma cell line, Yumm5.2, using a library focused on all metabolic genes (Zhu et al., 2021) and analyzed genes related to nucleotide synthesis, catabolism, and salvage, similar to those in human cells. In Yumm5.2 cells, using a different hairpin targeting *Cdkn2a* (**Fig. S1D**), almost all nucleotide metabolism genes dropped out (**Fig. 1C and Table S3**). Data taken from Dependency Map (DepMap) using cutaneous melanoma cell lines also indicated increased dependency on several nucleotide metabolism genes based on *CDKN2A* expression (**Fig. S1E**). Together, these screens identified 31 common genes in both nucleotide catabolism and biosynthesis pathways, including the PPP and one carbon metabolism (**Table S4**). These data suggest multiple nucleotide metabolism pathways are a vulnerability in cells with low p16/*CDKN2A* expression.

**Figure 1.**
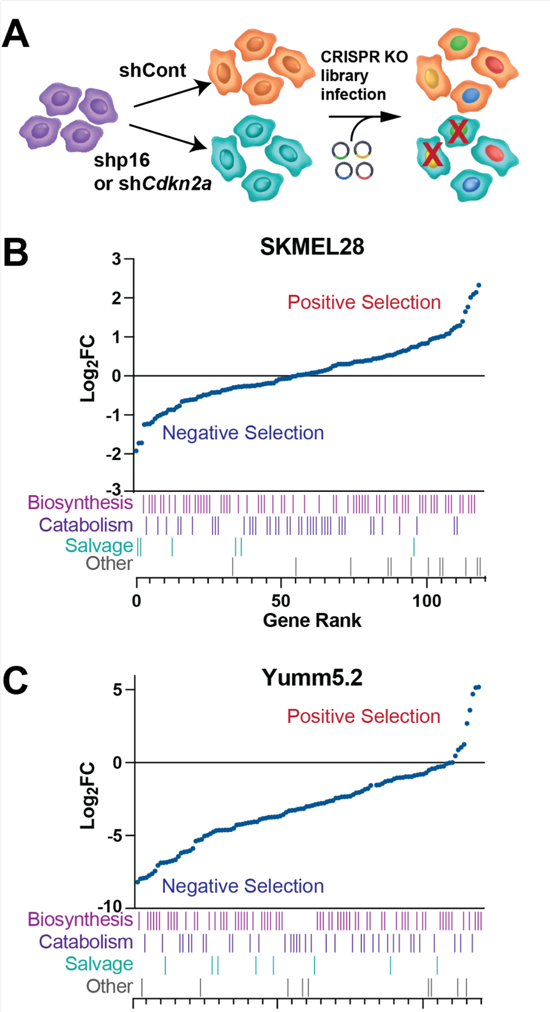
Multiple CRISPR knockout screens identify nucleotide metabolism genes that are selectively depleted in shp16/sh*Cdkn2a* cells. (A) Schematic of our CRISPR screens. Isogenic p16/*Cdkn2a* wildtype cells were infected with lentiviruses expressing shGFP control (shCont), shp16 (human), or sh*Cdkn2a* (mouse). These cells were infected with the CRISPR gRNA libraries at an MOI of 0.2-0.3. After 14 days in culture, gDNA was harvested and sequenced. **(B-C)** Analysis of 128 genes related to nucleotide synthesis, salvage, and catabolism identified multiple genes that are negatively selected in shp16/*shCdkn2a* vs. shControl in **(B)** human SKMEL28 and **(C)** mouse Yumm5.2 melanoma cells.

### p16/*CDKN2A* expression negatively correlates with nucleotide metabolism gene expression, polysome enrichment, and protein expression

Next, we cross-compared these 128 nucleotide metabolism genes to RNA-Seq data and found that multiple genes were both depleted in the CRISPR screen and transcriptionally upregulated in shp16 cells (Fig. 2A **and Table S5**). As we previously published that the PPP enzyme *RPIA* is translationally regulated (Buj et al., 2019), we also performed polysome fractionation followed by RNA-Seq (Poly-Seq). This analysis identified additional nucleotide metabolism transcripts that are also both depleted in the CRISPR screen and translationally upregulated in shp16 cells (Fig. 2B **and Table S6**). Of the 31 common genes, 23 (∼75%) are either transcriptionally or translationally upregulated in shp16 cells and many correspond to either *de novo* purine synthesis or one carbon metabolism (Fig. 2C). Using data from the DepMap, we found many of these 23 genes and proteins correlate with *CDKN2A* mRNA in cutaneous melanoma cell lines (Fig. 2D-E). Together, these data demonstrate upregulation of multiple nucleotide metabolism enzymes in the context of p16/*CDKN2A* loss.

**Figure 2.**
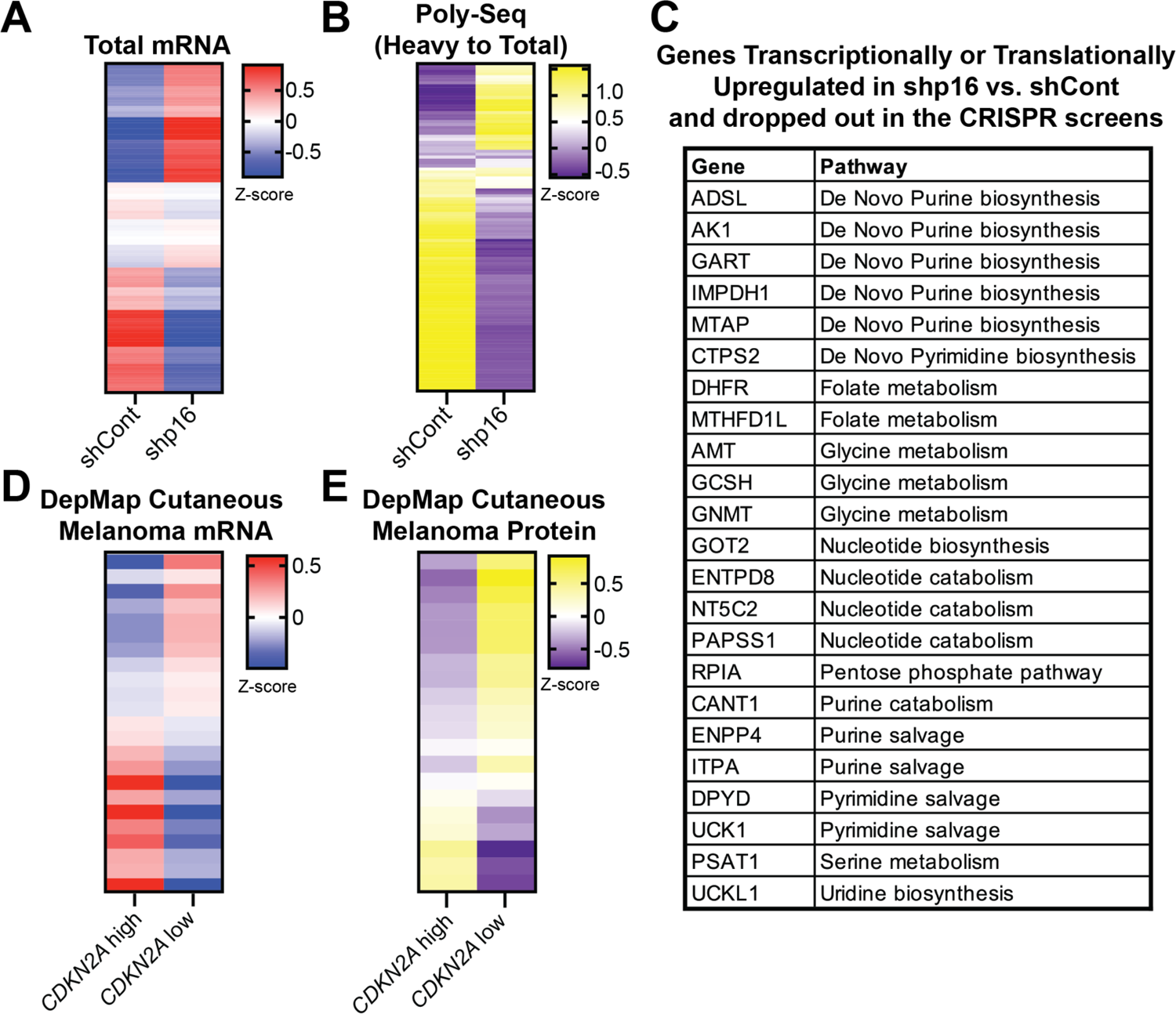
p16/*CDKN2A* negatively correlates with mRNA and protein expression of multiple nucleotide metabolism genes. **(A-B)** SKMEL28 human melanoma cells were infected with lentivirus expressing a short hairpin RNA (shRNA) targeting p16 (shp16). shGFP was used as a control (shCont). **(A)** Expression of the 128 nucleotide metabolism genes from RNA-Seq. **(B)** Polysome fractionation was performed and both heavy (>2 ribosomes) and total mRNA were sequenced, and the ratio of Heavy to Total was used to assess transcripts with increased translation. **(C)** Genes that are transcriptionally or translationally upregulated in shp16 SKMEL28 cells. **(D-E)** DepMap data of cutaneous melanoma cell lines. **(D)** mRNA expression of 31 genes identified in the CRISPR screens. **(E)** Protein expression of 31 genes identified in the CRISPR screens.

### Inhibition of *de novo* purine synthesis via multiple pathways decreases proliferation in p16/*Cdkn2a* knockdown cells

Our CRISPR dropout screens demonstrate that multiple nucleotide metabolism pathways may be a vulnerability of p16/*Cdkn2a* knockdown cells (Fig. 1), and further analysis indicates upregulation of *de novo* purine synthesis and one carbon metabolism genes in shp16 and *CDKN2A*-low cells (Fig. 2). We therefore used a variety of inhibitors that target these pathways (Fig. 3A). Notably, three enzymes targeted by these inhibitors were identified in our CRISPR screen (DHFR, IMDPH1, and GART). We determined that human isogenic cells with p16 knockdown have decreased proliferation upon treatment with many of these inhibitors *in vitro* (Fig. 3B). Additionally, knockdown or knockout of *Cdkn2a* in mouse cells decreased proliferation upon treatment with methotrexate, lometrexol, and aminopterin (**Fig. S2A-C**). We also found a correlation between *CDKN2A* expression and sensitivity to a variety of drugs that target nucleotide synthesis and antimetabolites in cutaneous melanoma cell lines on DepMap (Fig. 3C). Interestingly, while prior reports have demonstrated that *MTAP* deletions, which are near the same chromosomal locus as *CDKN2A*, increase sensitivity to *de novo* purine inhibitors, we did not observe decreased *MTAP* expression in our human or mouse knockdown cells (**Fig. S2D**). These experiments provide pharmacological evidence that p16/*Cdkn2a*-low cells are overall more sensitive to inhibition of nucleotide metabolism pathways, including *de novo* purine synthesis and folate synthesis, which corroborates our *in vitro* CRISPR screens (Fig. 1).

**Figure 3.**
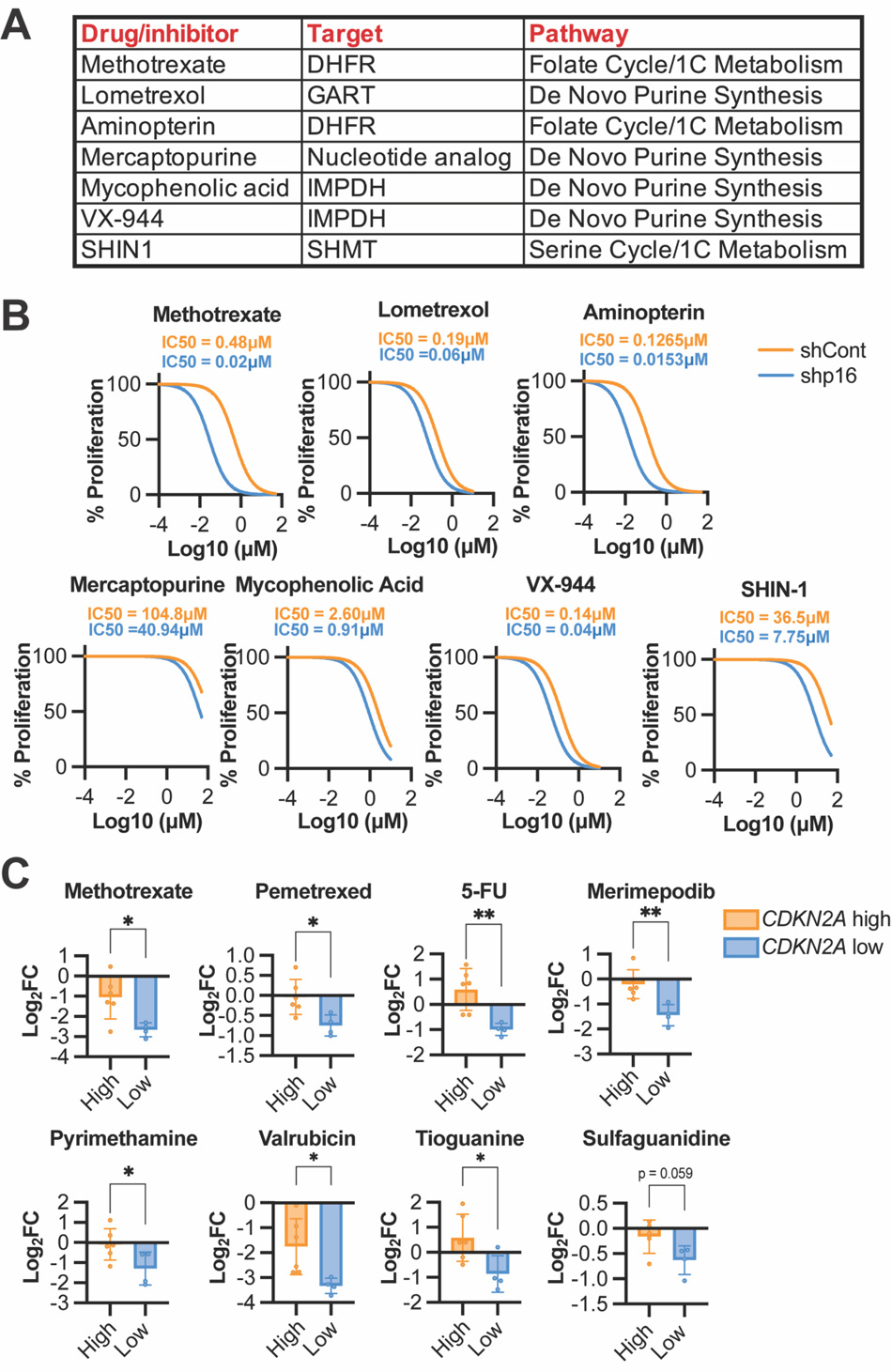
p16/*CDKN2A*-low cells are more sensitive to inhibitors of nucleotide metabolism. **(A)** Table of inhibitors used in *in vitro* cell line studies. 1C metabolism = one carbon metabolism. **(B)** SKMEL28 human melanoma cells were infected with lentivirus expressing a short hairpin RNA (shRNA) targeting p16 (shp16). shGFP was used as a control (shCont). Cells were treated with the indicated inhibitors and proliferation was assessed by crystal violet staining. One of 2-3 independent experimental replicates is shown (n=6). **(C)** Increased drug sensitivity from DepMap data of cutaneous melanoma cell lines with high or low *CDKN2A* expression. Data are mean ± SD. T-test. *p<0.05, **p<0.01

### Anti-folates induce apoptosis in p16/*Cdkn2a* knockdown cells

Next, we aimed to explore the mechanism of decreased proliferation in p16/*Cdkn2a* knockdown cells. We observed an increase in cytotoxicity in shp16/sh*Cdkn2a* cells treated with anti-folates compared to controls (Fig. 4A and **Fig. S3A**). Annexin V/PI staining was increased in shp16/sh*Cdkn2a* cells compared to controls, suggesting that these cells are undergoing apoptosis after treatment with these inhibitors (Fig. 4B **and Fig. S3B**), although the differences in Yumm5.2 cells were less robust than in SKMEL28. We did not observe an increase in senescence in shp16 cells treated with methotrexate (MTX) or lometrexol (LTX), as assessed by senescence-associated beta-galactosidase (SA-β-Gal) staining (**Fig. S3C**). This is in contrast to the CDK4/6 inhibitor palbociclib, which induced robust SA-β-Gal staining in shp16 cells (**Fig. S3D**), although the senescence-associated secretory phenotype (SASP) was blunted (**Fig. S3E**), consistent with our prior report that loss of p16 abrogates the SASP (Buj et al., 2021). While shp16 cell proliferation after MTX or LTX washout remained low, palbociclib-treated cells rebounded to near control levels (**Fig. S3F**). These data demonstrate that inhibition of *de novo* purine synthesis via the folate cycle induces apoptosis and also show that these effects are more long-term than inhibiting the CDK4/6 axis in p16-low cells. Many studies have reported a role for the SASP in the anti-tumor response to CDK4/6 inhibitors (Wagner and Gil); therefore, our data have multiple implications for the clinical use of these inhibitors in the context of p16/*CDKN2A* loss.

**Figure 4.**
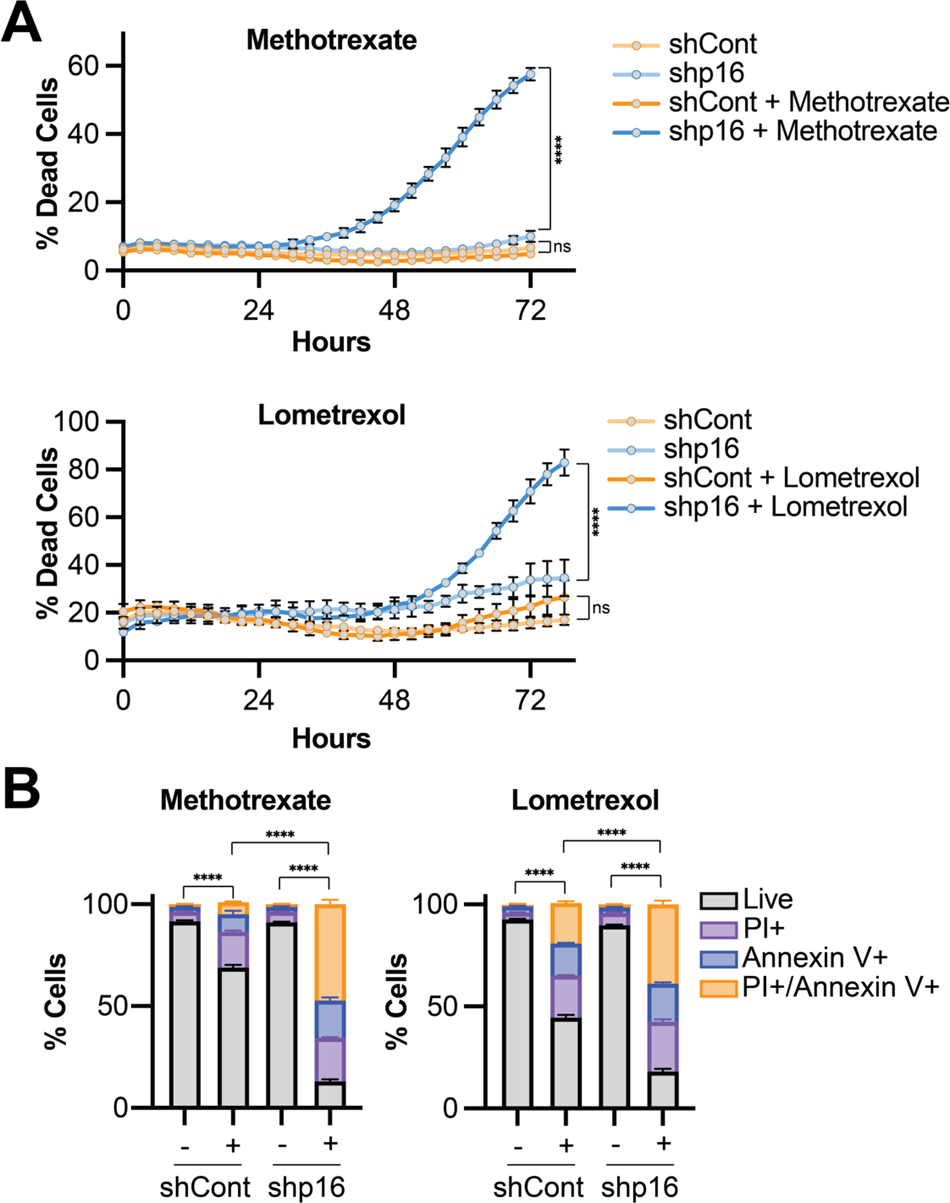
Multiple anti-folates induce apoptosis in p16 knockdown cells. **(A-B)** SKMEL28 human melanoma cells were infected with lentivirus expressing a short hairpin RNA (shRNA) targeting p16 (shp16). shGFP was used as a control (shCont). **(A)** Cells were treated with the indicated inhibitors and cytotoxicity was assessed using IncuCyte Cytotox Green reagent. One of 3 independent experimental replicates is shown (n=6). Data are mean ± SD. One-way ANOVA at endpoint. ****p<0.0001 **(B)** Cells were treated with the indicated inhibitors, and apoptosis was assessed using Annexin V/PI staining by flow cytometry. One of 3 independent experimental replicates is shown (n=3). Data are mean ± SD. One-way ANOVA of live cell population. ****p<0.0001

### Knockdown of p16 *in vivo* decreases tumor burden in response to the anti-folate methotrexate

We observed a decrease in proliferation and increase in death of shp16 cells treated with the anti-folate MTX (**Fig. 3-4**). Thus, we determined the effects of MTX in subcutaneous SKMEL28 tumors with and without p16 knockdown. While we observed large variability in tumor growth, there was a significant difference in tumor growth rate between shp16 tumors treated with MTX vs. vehicle controls (p=0.031) at day 30 when the first vehicle treated mice reached IACUC endpoints (1000mm^3^) and a trend towards increased survival in mice bearing shp16 tumor treated with methotrexate (Fig. 5A **and S4A**). Control tumors did not show appreciable differences in growth between treatment and vehicle control (p=0.734), and mice bearing control tumors did not have a significant difference in survival between vehicle and MTX treatment groups at the time the shp16 tumors all reached endpoint (Fig. 5B and S4B). Interestingly, comparison between control tumors and shp16 MTX treated tumors was highly significant both at Day 30 (p=0.002) and at the timepoint when all shp16 mice had reach endpoint (Day 55, p<0.001). MTX is known to have side effects, including decreased white blood cell counts (Hansen Hh Fau - Selawry et al.). We did not observe differences in body weight or blood cell counts in mice bearing shp16 tumors treated with MTX at the dose and duration used (**Fig. S4C-D**). These data provide evidence that MTX may be clinically beneficial for tumors with low p16 expression, although additional experiments with refined dosing and in other models are needed.

**Figure 5.**
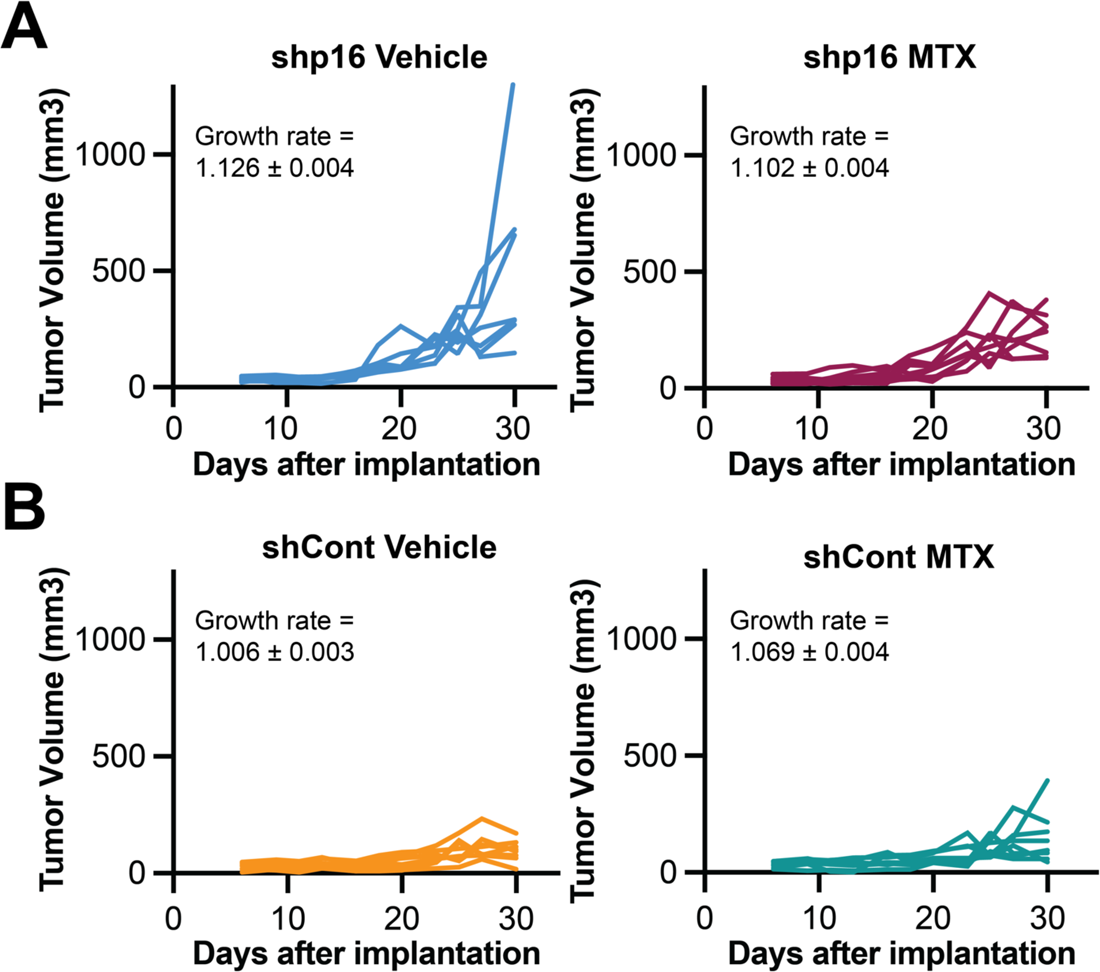
*In vivo* shp16 tumors are sensitive to the anti-folate methotrexate. **(A-B)** SKMEL28 human melanoma cells were infected with lentivirus expressing a short hairpin RNA (shRNA) targeting GFP (shCont) or p16 (shp16). 10^7^ cells were subcutaneously implanted into athymic nude mice. Individual tumor growth curves in the indicated groups. Shown are growth rates ± SE. Linear mixed-model group comparisons: shp16 vs. shCont p<0.001; shCont: MTX vs. Vehicle p=0.734; shp16: MTX vs. Vehicle p=0.031; (shp16: MTX vs. Vehicle) vs. (Control: Vehicle vs MTX) p=0.002.

### *De novo* purine synthesis and one carbon metabolism genes correspond to worse melanoma patient survival

Finally, using TCGA data, we found that upregulation of *de novo* purine synthesis and one carbon metabolism genes occurs in many metastatic melanomas, although *MTAP* is typically downregulated due to deletion of chromosome 9p21 (Zhang et al., 1996) (Fig. 6A and **Table S7**). Moreover, alteration of these genes corresponds with worse overall survival (Fig. 6B), whereas upregulation of genes involved in *de novo* pyrimidine synthesis, overall nucleotide biosynthesis, or nucleotide salvage do not have an association with melanoma patient survival (Fig. 6C-E). Together, these data show that alterations in genes related to *de novo* purine synthesis and one carbon metabolism are specifically associated with melanoma patient survival.

**Figure 6.**
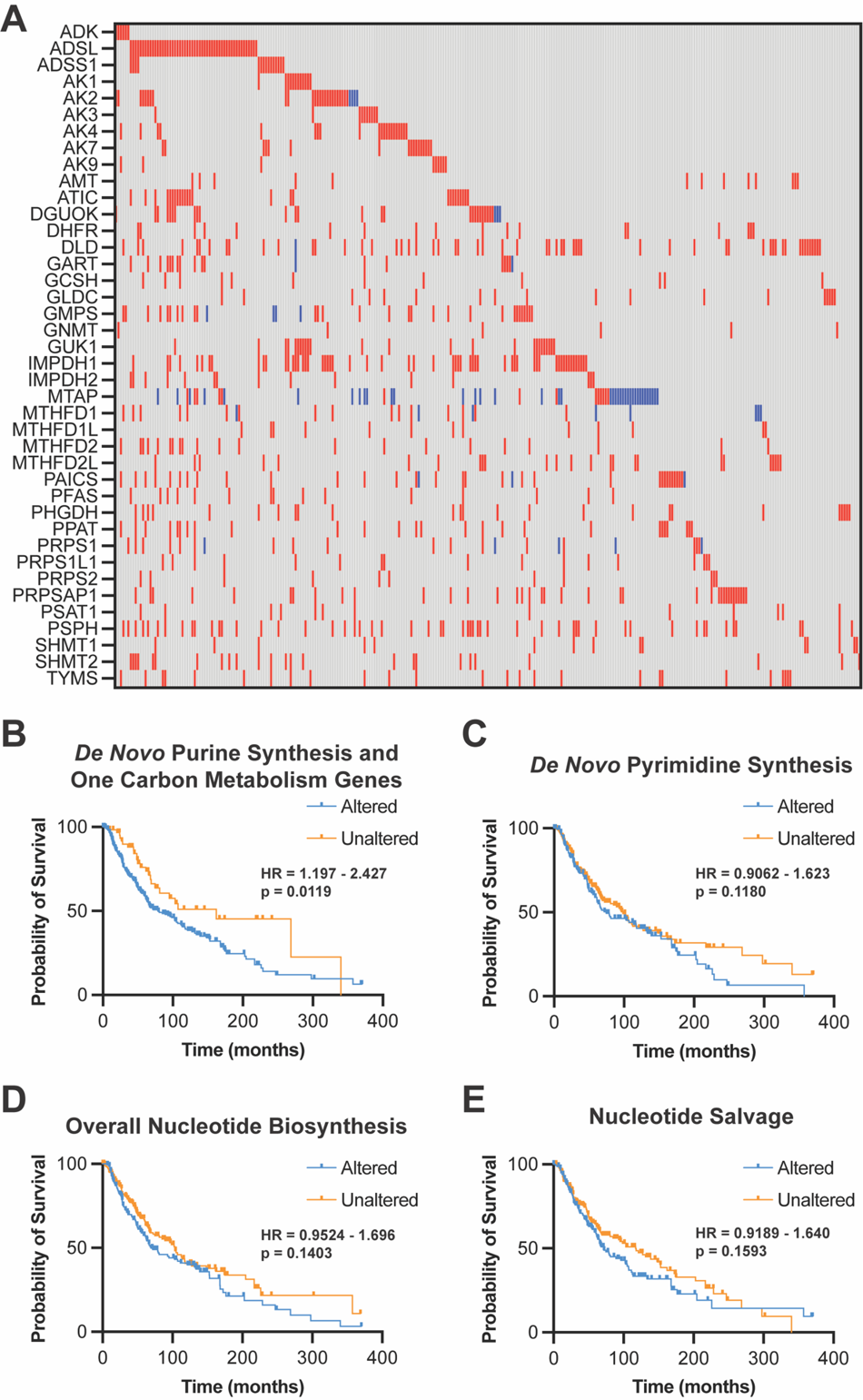
*De novo* purine synthesis and one carbon metabolism genes are associated with worse overall survival in metastatic melanomas. Data from The Cancer Genome Atlas (TCGA) Skin Cutaneous Melanoma PanCancer Atlas (367 metastatic melanomas). (A) *De novo* purine synthesis and one carbon metabolism genes in TCGA metastatic melanoma samples. Red indicates increased mRNA expression. Blue indicates decreased mRNA expression. Overall survival probability of patients with alterations in *de novo* purine synthesis and one carbon metabolism genes **(B)**, *de novo* pyrimidine genes **(C)**, overall nucleotide biosynthesis genes **(D)**, and nucleotide salvage genes **(E)**. Log rank p value and 95% confidence interval.

## Discussion

We previously found that p16 loss increases nucleotide synthesis via mTORC1 and the PPP (Buj et al., 2019). Here we identified multiple other nucleotide metabolic genes that are potential vulnerabilities of p16-low cells. Upon direct comparison of multiple anti-folates and anti-metabolites in both human and mouse p16 wildtype and shp16/sh*Cdkn2a* cells, we found that p16 status corresponds to therapeutic response to many of these agents. These agents induce cell death, unlike the CDK4/6 inhibitor palbociclib that induces senescence. Given the context-dependent effects of senescence on the microenvironment (detailed below), these data suggest that *de novo* purine synthesis or one carbon metabolism inhibitors may provide therapeutic benefit for the ∼50% of human tumors with low p16/*CDKN2A* expression.

Previous studies and clinical trials found that most chemotherapies have little clinical benefit in melanoma patients (Luke and Schwartz, 2013). However, to date, trials have not stratified patients based on p16/*CDKN2A* expression. Our data suggest that while patients with wildtype p16/*CDKN2A* or high to moderate p16/*CDKN2A* expression would likely not robustly respond to these therapies, those with low or mutant p16/*CDKN2A* may benefit. Moreover, the *CDKN2A* gene is located on chromosome 9p21 and is often deleted in tandem with other genes in that locus including *MTAP* (Zhang et al., 1996). Prior work has found that patients with *MTAP* deletions are more responsive to *de novo* purine synthesis inhibitors, including anti-folates (Alhalabi et al., 2022; Hu et al., 2021; Tang et al., 2018), although we did not observe a decrease in *MTAP* upon p16 knockdown (**Fig. S2D**). Thus, cancers with homozygous deletion of the entire locus containing both genes would presumably be highly sensitive to these inhibitors, although this will need to be tested experimentally in isogenic cells. Additionally, we previously demonstrated that p16 loss increases mTORC1 activity (Buj et al., 2019), and a recent study found that cancers with increased mTORC1, such as those with TSC2 deficiency, are highly sensitive to inhibitors of IMPDH (Valvezan et al., 2020). Together, these data open up new questions regarding the importance of stratification of patients for future anti-metabolite therapy based on p16/*CDKN2A* status.

Multiple anti-folates increased cell death in shp16/sh*Cdkn2a* cells, although the mechanism remains to be determined. p16 knockdown cells proliferate faster than p16 wildtype cells (**Fig. S1B**). Increased proliferation would increase the demand for dNTPs and likely other nucleotides required for biomass. Indeed, many chemotherapies are more effective in highly proliferative cells, and the status of other cell cycle regulators such as RB1 and CDK12 affect response to antifolates and anti-metabolites (Derenzini et al., 2008; Filippone et al., 2022; Robinson et al., 2013). We cannot rule out other effects of these agents, such as redox imbalance due to inhibition of glutathione synthesis or NADPH, changes in methylation due to methionine cycle dysregulation, or energy stress due to decreased ATP. Indeed, one paper has linked p16 deficiency to increased mitochondrial ROS that is uncoupled from cell cycle regulation (Tyagi et al., 2017), again suggesting a need to better understand non-canonical roles of p16 when mechanistically investigating therapies. Another recent paper found that imbalanced nucleotides decrease proliferation (Diehl et al., 2022); thus, it is also possible that the anti-metabolites used here create a nucleotide imbalance that perturb S phase. Future work will be required to more fully elucidate the mechanisms of sensitivity, which may provide additional evidence into the metabolic reprogramming induced by p16/*CDKN2A* loss. Interestingly, recent reports have demonstrated that media and FBS conditions are critically important for response to MTX (Abbott et al., 2023; Kyle et al., 2023). Future studies are therefore also needed to determine whether the differential sensitivity observed is recapitulated in more physiological conditions.

Melanoma standard-of-care is now immunotherapy (Larkin et al., 2015; Robert et al., 2015a; Robert et al., 2015b), although >40% of patients do not achieve a significant response (Callahan et al., 2018; Robert et al., 2015b; Valsecchi, 2015; Wolchok et al., 2017). Interestingly, recent data demonstrate that a decreased purine to pyrimidine ratio leads to a nucleotide imbalance that elevates the expression of the immunoproteasome and enhances anti-PD1 response (Keshet et al., 2020; Lee et al., 2018). Another study found that the anti-folate pemetrexed induces immunogenic cell death and augments T cell function to synergize with anti-PD1 (Schaer et al., 2019). Therefore, these inhibitors may promote an anti-tumor immune response that could be exploited in combination with immunotherapy. Indeed, multiple trials are ongoing to better understand whether these agents can be used in combination in a variety of solid tumors, including melanoma (Lelliott et al., 2022) (clinicaltrials.gov). While these drugs are also potent immunosuppressants, our data suggest there is a therapeutic window in p16/*CDKN2A*-low tumors that may allow for an antitumor response that does not have marked effects on white blood cell counts (**Fig. S4E**), although additional experiments are needed.

CDK4/6 inhibitors are currently in clinical trials in combination with immunotherapies in melanoma and other solid tumors (Ettl et al., 2022; Lelliott et al., 2022; Liu et al., 2020; Moore et al., 2022) (clinicaltrials.gov). These inhibitors induce cytostasis, generally associated with a senescence arrest (Klein et al., 2018). Indeed, we observed increased SA-β-Gal activity in shp16 cells treated with the CDK4/6 inhibitor palbociclib (**Fig. S3D**). However, cells began to proliferate again after washout of palbociclib, suggesting that this is either a pseudosenescent phenotype or a population of cells that do not senesce are capable of repopulating after discontinuing therapy. Induction of senescence may promote an anti-tumor immune response via the SASP (Chen et al., 2023; Hao et al., 2021; Marin et al., 2023; Paffenholz et al., 2022; Xue et al., 2007), although other studies suggest senescence and the SASP is immunosuppressive (Faget et al., 2019; Oesterreich and Aird, 2023; Shahbandi et al., 2022). We previously published that loss of p16/*CDKN2A* expression corresponds to decreased SASP (Buj et al., 2021), and our data here indicate a blunted SASP induction in p16 knockdown cells treated with palbociclib (**Fig. S3E**). This suggests that while CDK4/6 inhibitors induce a potent cytostatic effect in p16-low cells, they may not have the added benefit to potentiate immunotherapy due to suppressed pro-inflammatory factors. Experiments to determine the extent of synergy between CDK4/6 inhibitors and immunotherapy in the context of p16/*CKDN2A* expression will be helpful towards understanding this combination.

In summary, we identified multiple metabolic vulnerabilities of p16/*CDKN2A*-low cancer cells that can be exploited using various anti-metabolites or anti-folates. While we focused on melanoma cells in this study, these results may have implications for other human tumors where p16/*CDKN2A* expression is lost as we have previously published that metabolic changes due to p16 loss are a universal phenomenon (Buj et al., 2019). Low expression of p16/*CDKN2A* may open up a therapeutic window for these agents that kill tumor cells, although this will need to be further investigated in humans.

## Supplemental Figures

**Figure S1.**
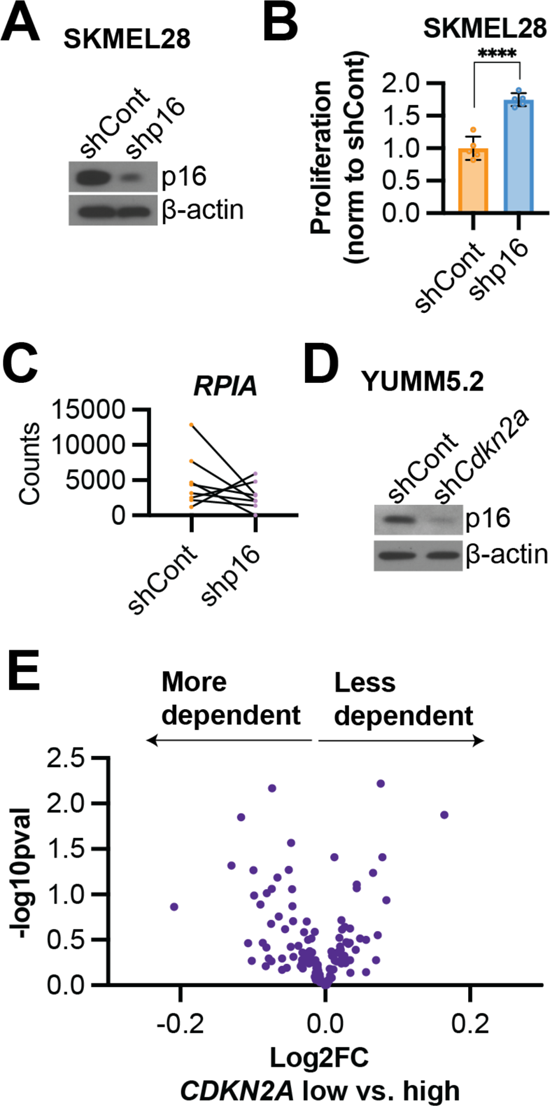
Knockdown of p16/*Cdkn2a* in human and mouse melanoma cell lines and DepMap dependency score data based on *CDKN2A* expression. Related to Figure 1. **(A-B)** SKMEL28 human melanoma cells were infected with lentivirus expressing a short hairpin RNA (shRNA) targeting p16 (shp16). shGFP was used as a control (shCont). **(A)** Immunoblot analysis of p16. β-actin was used as a loading control. One of 6 independent experimental replicates is shown. **(B)** Proliferation was assessed by crystal violet staining. One of 6 independent experimental replicates is shown (n=6). Data are mean ± SD. T-test. ****p<0.001 **(C)** Counts for each *RPIA* gRNA in the SKMEL28 CRISPR KO screen (**see** Fig. 1B). **(D)** Yumm5.2 mouse melanoma cells were infected with lentivirus expressing a shRNA targeting *Cdkn2a* (sh*Cdkn2a*). shGFP was used as a control (shCont). Immunoblot analysis of p16. β-actin was used as a loading control. One of 3 independent experimental replicates is shown. **(E)** Volcano plot of dependency data from DepMap of cutaneous melanoma cell lines based on *CDKN2A* expression.

**Figure S2.**
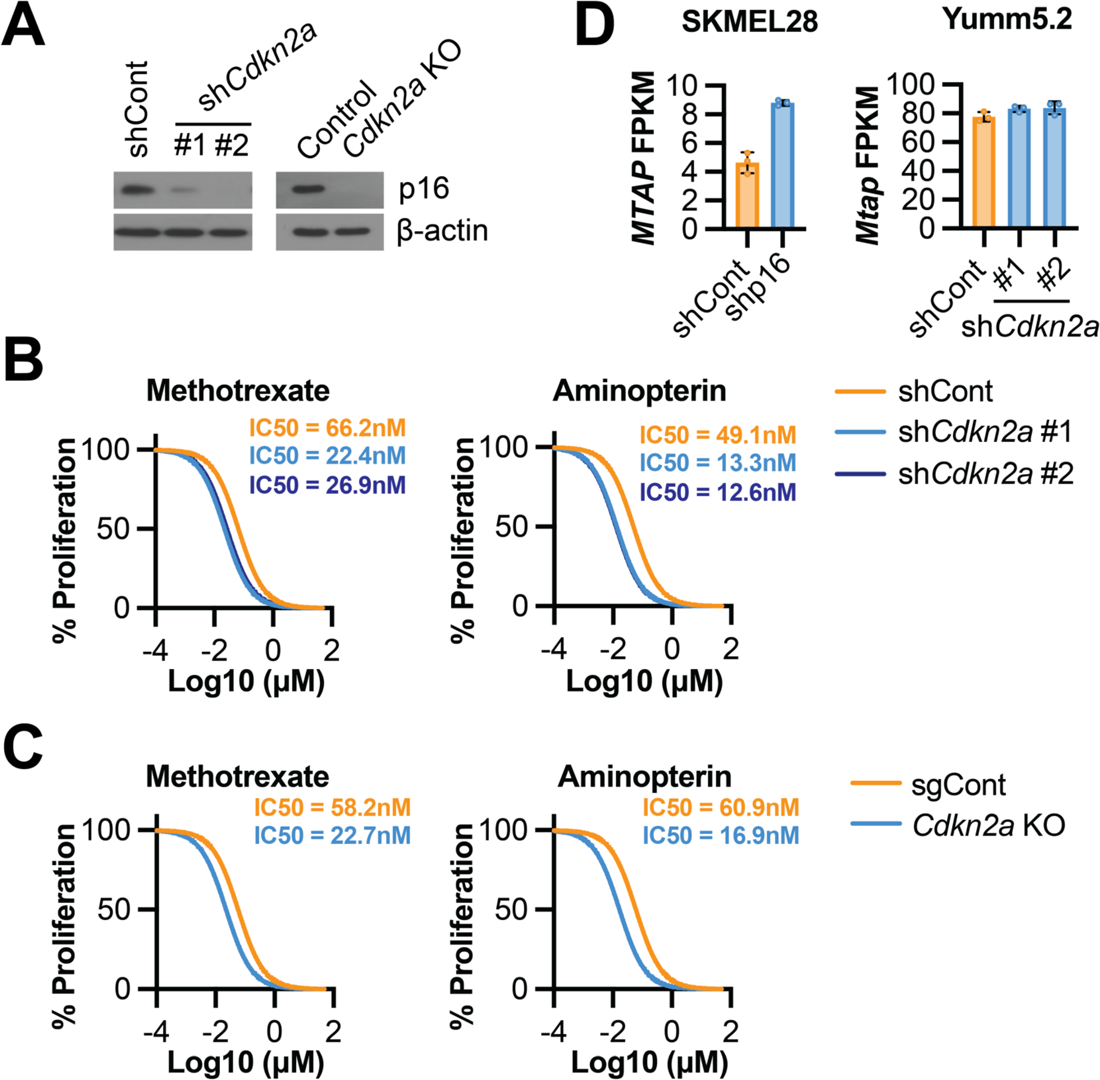
Knockdown or knockout of *Cdkn2a* in mouse melanoma cell lines increases sensitivity to multiple anti-folates. Related to Figure 3. **(A-B)** Yumm5.2 mouse melanoma cells were infected with lentivirus expressing short hairpin RNAs (shRNA) targeting *Cdkn2a* (sh*Cdkn2a* #1 and #2) or a gRNA targeting *Cdkn2a*. shGFP was used as a control for KD experiments (shCont). A gRNA targeting an intergenic region was used as a control for KO experiments. **(A)** Immunoblot analysis of p16. β-actin was used as a loading control. One of 3 independent experimental replicates is shown. **(B-C)** Cells were treated with the indicated inhibitors, and proliferation was assessed by crystal violet staining. One of 2-3 independent experimental replicates is shown (n=3). (D) *MTAP* Fragments Per Kilobase of transcript per Million mapped reads (FPKM) in SKMEL28 human melanoma cells with p16 knockdown (shp16) and Yumm5.2 mouse melanoma cells with *Cdkn2a* knockdown (sh*Cdkn2a* #1 and #2). Data are mean ± SD.

**Figure S3.**
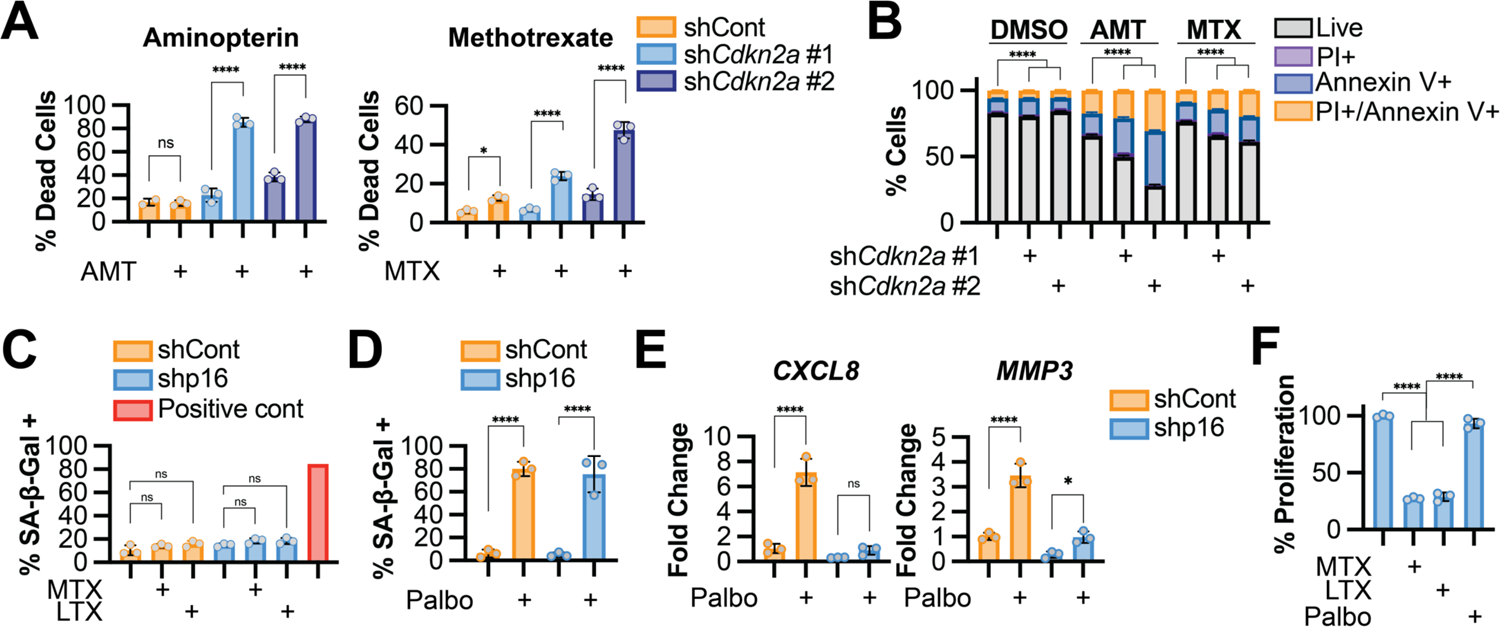
Anti-folates induce death in sh*Cdkn2a* cells; shp16 cells do not mount a robust senescence-associated secretory phenotype (SASP) in response to Palbociclib; and antiproliferative effects of anti-folates are longer-lived than palbociclib. Related to Figure 4. **(A-B)** Yumm5.2 mouse melanoma cells were infected with lentivirus expressing short hairpin RNAs (shRNA) targeting *Cdkn2a* (sh*Cdkn2a* #1 and #2). **(A)** Cells were treated with the indicated inhibitors and cytotoxicity was assessed using IncuCyte Cytotox reagent. One of 2 independent experimental replicates is shown (n=3). **(B)** Cells were treated with the indicated inhibitors, and apoptosis was assessed using Annexin V/PI staining by flow cytometry. One of 2 independent experimental replicates is shown (n=3). Statistical analysis of live cells is shown. **(C-F)** SKMEL28 human melanoma cells were infected with lentivirus expressing a short hairpin RNA (shRNA) targeting p16 (shp16). shGFP was used as a control (shCont). **(C)** Cells were treated with the indicated inhibitors, and senescence-associated beta-galactosidase (SA-β-Gal) activity was assessed. Cisplatin was used as a positive control. One of 3 independent experimental replicates is shown (n=3). **(D)** Cells were treated with 1µM palbociclib, and SA-β-Gal activity was quantified. One of 3 independent experimental replicates is shown (n=3). **(E)** SASP gene expression. One of 3 independent experimental replicates is shown (n=3). **(F)** Cells were treated with the indicated drugs for 4 days, after which the drugs were washed out. After an additional 5 days in culture, proliferation was assessed by crystal violet staining. One of 2 independent experimental replicates is shown (n=3). Data are mean ± SD. One-way ANONA. *p<0.05; ****p<0.001; ns = not significant. AMT: aminopterin; MTX: methotrexate; LTX: lomotrexol; palbo: palbociclib

**Figure S4.**
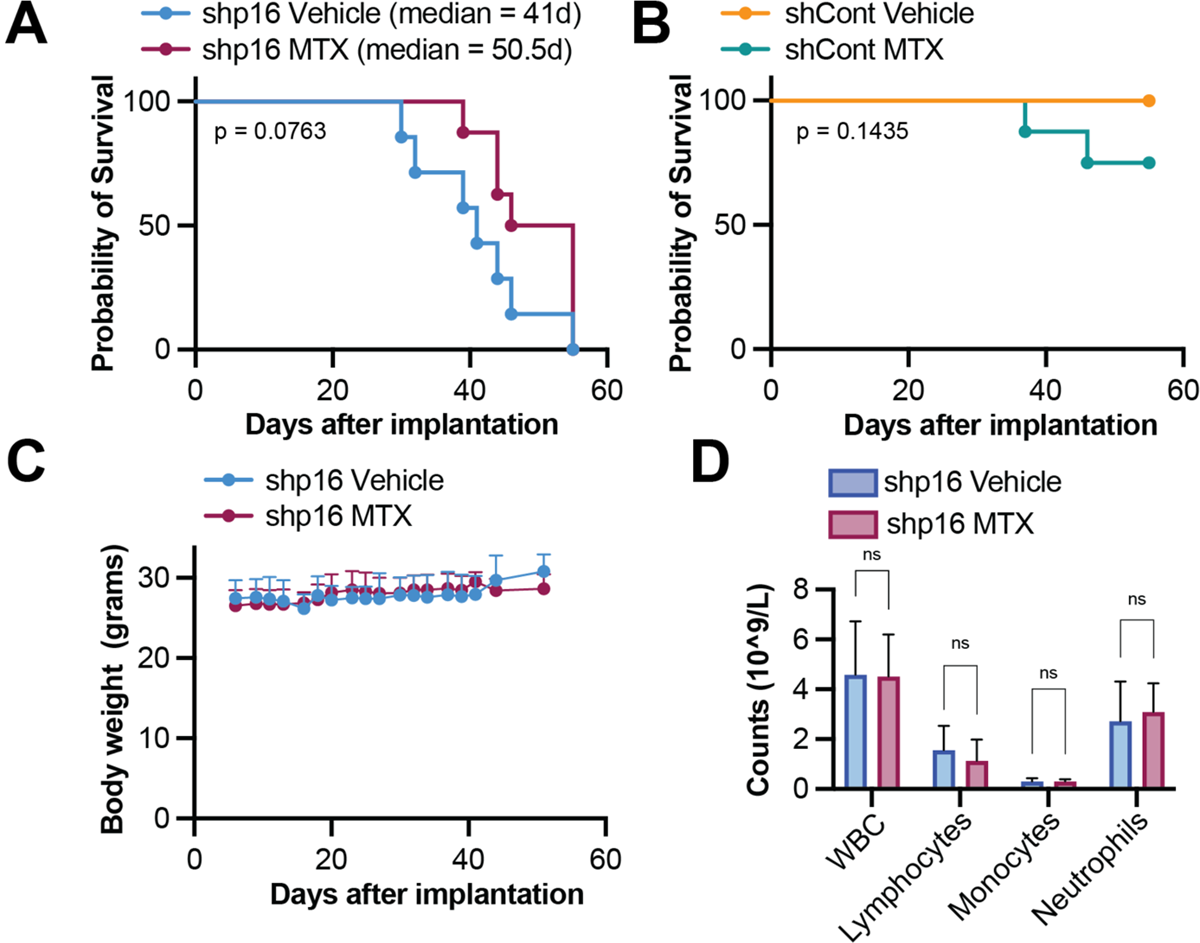
shp16 tumor bearing mice treated with methotrexate have a trend towards a survival advantage; and methotrexate does not affect body weight or white blood cell counts. Related to Figure 5. **(A-D)** SKMEL28 human melanoma cells were infected with lentivirus expressing a short hairpin RNA (shRNA) targeting GFP (shCont) or p16 (shp16). 10^7^ cells were subcutaneously implanted into athymic nude mice. **(A-B)** Kaplan-Meier survival curves at the timepoint when all shp16 tumor bearing mice reached endpoint (1000mm^3^). Log-rank p values are shown. **(C)** Mice body weight over time. **(D)** Blood cell counts of the indicated shp16 mice at endpoint. Data are mean ± SD. T-test. ns = not significant

## Acknowledgements

We would like to thank Fran Vazquez for help with the CRISPR schematic. This work was supported by grants from the National Institutes of Health (R37CA240625 to K.M.A., R01CA259111 to K.M.A. and N.W.S, and T32GM133332 to A.R.C.), the Melanoma Research Foundation (to R.B.G.), and the Ovarian Cancer Research Alliance (MIG-2023-2-1018 to A.U.). This project used the following Hillman Cancer Center Flow Facility, Cancer Bioinformatics Services, Cancer Biostatistics Services, Cancer Genomics Facility, and Animal Facility shared resources which are supported in part by award P30CA047904.

## Author Contributions

**Naveen Kumar Tangudu:** Investigation, Methodology, Visualization, Writing – Review & Editing. **Raquel Buj:** Investigation, Visualization, Writing – Review & Editing. **Hui Wang:** Investigation. **Jiefei Wang:** Software, Formal Analysis. **Aidan Cole:** Investigation, Writing – Review & Editing. **Apoorva Uboveja:** Investigation, Writing – Review & Editing. **Richard Fang:** Investigation, Writing – Review & Editing. **Amandine Amalric:** Investigation, Writing – Review & Editing. **Peter Sajjakulnukit: Investigation. Maureen A. Lyons:** Methodology. **Kristine Cooper:** Statistical analysis. **Nadine Hempel:** Writing – Review & Editing**. Nathaniel W. Snyder:** Writing – Review & Editing, Funding Acquisition**. Costas A. Lyssiotis:** Methodology, Supervision. **Uma R. Chandran:** Methodology, Supervision. **Katherine M. Aird:** Conceptualization, Visualization, Writing – Original Draft, Writing – Review & Editing, Supervision, Project Administration, Funding Acquisition.

## Materials and Methods

### Cell Lines

SKMEL28 and YUMM5.2 cells were purchased from ATCC. SKMEL28 cells were cultured in DMEM (Fisher Scientific cat# MT10013CV,) supplemented with 5% Fetal Bovine Serum (BioWest, cat# S1620). YUMM5.2 cells were cultured in DMEM (Fisher Scientific, cat# MT10092CV,) supplemented with 10% Fetal Bovine Serum (BioWest, cat# S1620) and nonessential amino acids (Corning, cat#25-025-Cl). Both were supplemented with 1% Penicillin/Streptomycin (Fisher Scientific, cat#15-140-122). All cell lines were routinely tested for mycoplasma as described in (Uphoff and Drexler, 2005).

### Lentiviral packaging and infection

The human shRNA hairpin pLKO.1-shp16 (TRCN0000010482), the murine shRNA hairpins pLKO.1-shCdkn2a #1 (TRCN0000077816) and #2 (TRCN0000362595) and the human and murine pLKO.1-shGFP control (Addgene, cat#30323) vectors were packaged using the ViraPower Kit (Invitrogen, cat#K497500) following the manufacturer’s instructions. Cells were infected with corresponding vectors for 16h and selected for 3 days with 1µg/mL of puromycin (SKMEL28) or 3µg/mL puromycin (YUMM5.2).

### Design of *Cdkn2a* CRISPR KO murine cell line

Single-stranded oligonucleotides targeting *CDKN2A* (forward: CACCGGCTG-GATGTGCGCGATGCC and reverse: AAACGGCATCGCGCACATCCAGCC, Integrated DNA Technologies) or intergenic region control (forward: CACCGAGTGTTCCTAGAGATAGAAG and reverse: AAACCTTCTATCTCTCTAGGAACACTC) were annealed, phosphorylated, and ligated into pLentiCRISPRv2 (Addgene, cat#52961), kindly gifted by Feng Zhang. Lentivirus was packaged using the ViraPower Kit as described above and used to infect YUMM5.2 cells. Mixed pooled population YUMM5.2 cells infected with intergenic region control gRNAs were used as control while a single clone was obtained for YUMM5.2 cells targeted with *CDKN2A* gRNAs.

### Nucleotide Metabolic CRISPR library construction

We constructed a pooled sgRNA library containing 2800 sgRNAs targeting various genes either directly or indirectly related to nucleotide metabolism in addition to controls targeting intragenic regions as described previously (Joung et al., 2017). Briefly, we used publicly available CRISPR sgRNA design tools that optimize on-target and minimize off-target genome editing (http://crispr.dfci.harvard.edu/SSC/) and pooled human metabolic library (Birsoy et al., 2015) to identify 10 sgRNAs for each gene. The pooled oligo library was synthesized by Twist Bioscience. To the oligo library was cloned into lentiCRISPRv2, Addgene cat#52961) by PCR amplification using NEB Next High-Fidelity PCR Master Mix (New England Biolabs, cat#M0541S). The PCR product was digested with Esp3I (BsmBI) (Fisher Scientific, cat#FERFD0454) and ligated into the cut lentiCRISPRv2 backbone. The library was sequenced prior to its use to confirm coverage.

### CRISPR-based Screens

The human nucleotide-focused CRISPR KO library containing 128 genes involved in nucleotide biosynthesis, salvage, and catabolism (Table S1), was designed as stated above. The mouse metabolic-focused CRISPR KO library was a gift from Dr. Kivanc Birsoy (Addgene, cat#160129). The screening using the human nucleotide-focused library was conducted on SKMEL28 shGFP control and shp16 cells, while the screening using the mouse metabolic-focused library was conducted on YUMM5.2 shGFP control and sh*Cdkn2a* #1 cells. Briefly, the appropriate number of cells were infected with pooled libraries at an MOI <0.3 to achieve >400-fold library coverage after selection with puromycin. Cells were passaged every 2-3 days for two weeks maintaining the whole population. At the end of the experiments, cells were harvested for genomic DNA extraction using the Zymo Research kit (cat# D4069). sgRNA inserts were PCR amplified using Ex Taq DNA Polymerase (Takara, cat#RR001A) from sufficient genome equivalents of DNA to achieve an average coverage of >200x of the sgRNA library. See primers in **Table S8**. Pooled PCR amplicons were then sequenced. MAGeCK was used as the bioinformatics pipeline to analyze negatively and positively enriched genes (Li et al., 2014).

### Western blotting

Cells lysates were collected in 1X sample buffer (2% SDS, 10% glycerol, 0.01% bromophenol blue, 62.5mM Tris, pH 6.8, 0.1M DTT) and boiled to 95°C for 10 min. Protein concentration was determined using the Bradford assay (Bio-Rad, cat#5000006). An equal amount of total protein was resolved using SDS-PAGE gels and transferred to nitrocellulose membranes (Fisher Scientific) at 110mA for 2h at 4°C. Membranes were blocked with 5% nonfat milk or 4% BSA in TBS containing 0.1% Tween-20 (TBS-T) for 1 h at room temperature. Membranes were incubated overnight at 4°C in primary antibodies in 4% BSA/TBS + 0.025% sodium azide. Membranes were washed 4 times in TBS-T for 5 min at room temperature after which they were incubated with HRP-conjugated secondary antibodies (Cell Signaling Technology) for 1 h at room temperature. After washing 4 times in TBS-T for 5 min at room temperature, proteins were visualized on film after incubation with SuperSignal West Pico PLUS Chemiluminescent Substrate (ThermoFisher, Waltham, MA). Primary antibodies: Rabbit anti-CDKN2A (Abcam, ab108349; 1:1000); Rabbit anti-p16 INK4A (Cell Signaling Technology, 29271; 1:1000); Rabbit anti-CDKN2A/p19 ARF (Abcam, ab80; 1:1000); mouse anti-β-actin (Sigma Aldrich, A1978; 1:10000). Secondary antibodies: Antimouse IgG, HRP-linked (Cell Signaling Technology, 7076; 1:10,000), Anti-Rabbit IgG, HRP-linked (Cell Signaling Technology, 7074; 1:5000).

### Proliferation assays

An equal number of cells were seeded in multiwell plates and cultured for 4-5 days. Proliferation was assessed by fixing the plates for 5 min with 1% paraformaldehyde after which they were stained with 0.05% crystal violet. Wells were destained using 10% acetic acid. Absorbance (590nm) was measured using a spectrophotometer (BioTek Epoch Microplate reader).

### RNA isolation, Sequencing, and Analysis

Total RNA was extracted from cells with Triol (Ambion, cat#15596018) and DNase treated, cleaned and concentrated using Zymo columns (Zymo Research, cat#R1013) following manufacturer’s instructions. RNA integrity number (RIN) was measured using BioAnalyzer (Agilent Technologies) RNA 6000 Nano Kit to confirm RIN above 7 for each sample. The cDNA libraries, next generation sequencing, and bioinformatics analysis was performed by Novogene. Raw and processed RNA-Seq data can be found on GEO (GSE243717).

### Polysome Profiling and Sequencing

Eight culture plates per condition (∼23 million cells per condition) were incubated with harringtonine (2μg/mL) for 2 minutes at 37°C followed by 5 minutes of cycloheximide (100μg/mL) treatment at 37°C. Cells were washed twice with PBS after each treatment. Cells were scraped in 600uL of lysis buffer (50mM HEPES, 75mM KCl, 5mM MgCl2, 250mM sucrose, 0.1mg/mL cycloheximide, 2mM DTT, 1% Triton X-100 and 1.3% sodium deoxycholate and 5μl of RNase OUT) on ice. Lysates were rocked for 10 minutes at 4°C and centrifuged at 3000g for 15 minutes at 4°C. 400μl of lysates supernatant (cytosolic cell extracts) were layered over cold sucrose gradients (10mM HEPES, 75mM KCl, 5mM MgCl2, 0.5mM EDTA and increasing sucrose concentrations from 20% to 47%). Gradients were centrifuged at 34,000 rpms in a Beckman SW41 rotor for 2h and 40 minutes at 4°C. After centrifugation, low (0 to 2 ribosomes) and high (>2 ribosomes) polysome fractions were collected in Trizol (1:1) using a density gradient fractionation system (Brandel) equipped with a UA-6 absorbance detector and a R1 fraction collector. RNA was DNase treated, cleaned, and concentrated using Zymo columns (Zymo Research, cat# R1013). The cDNA libraries, next generation sequencing, and bioinformatics analysis was performed by Novogene. Raw and processed Poly-Seq data can be found on GEO (GSE243717).

### Dependency Map Data

For analysis of dependency scores, CRISPR (DepMap Public 23Q2+Score, Chronos) was downloaded for cutaneous melanoma cell lines. Cells were characterized as *CDKN2A* high or low based on mRNA expression (Expression Public 23Q2). Cell lines with known *CDKN2A* mutations were excluded. A similar method was used to analyze proteomics and drug sensitivity (using the PRISM Repurposing Primary Screen).

### IncuCyte cytotoxicity assay

Live cell viability was assessed using IncuCyte S3 imaging system (Sartorius). Briefly, cells were seeded in 96 well plate and treated with drug, either methotrexate (13960; Cayman Chemical Company) or lometrexol hydrate (18049; Cayman Chemical Company), in the presence of 1x Cytotox reagent (cat# ESS4633 or ESS4632; Sartorius), a highly sensitive cyanine nucleic acid dye. Cells were imaged for 3 days with live cell imaging every 3h point. Dead cell quantification was performed using the IncuCyte software.

### Annexin V/propidium iodide staining and flow cytometry

For Annexin V/7AAD experiments, cells (4 × 10^4^/well in 6-well plates) were treated for 5 days with 0.07μM methotrexate or 0.09μM lometrexol in DMEM + 5% FBS. Cells were trypsinized, counted in hemocytometer, and each FACS tube contained approximately 2 × 10^5^ harvested cells. Cells were stained with Annexin V (R37176; Thermo Fisher Scientific) in 2.5 mM Ca2+ containing DMEM for 15 min at room temperature. Prior to FACS analysis, 5µl (40ng/sample) propidium iodide added to cell suspension (eBiosciences, cat#00-6990-50). Data were analyzed using FlowJo software. Single stain controls were used for compensation.

### Senescence Associated-β-Galactosidase assay

Cells were treated with 1μM palbociclib (Sigma Aldrich, cat#PZ0383) for 5 days. SA-β-Gal staining was performed as previously described (Dimri et al., 1995). Cells were fixed in 2% formaldehyde/0.2% glutaraldehyde in PBS (5 min) and stained (40 mM Na_2_HPO_4_, 150 mM NaCl, 2 mM MgCl_2_, 5 mM K_3_Fe(CN)_6_, 5 mM K_4_Fe(CN)_6_, and 1 mg/ml X-gal) overnight at 37°C in a non-CO2 incubator. Images were acquired at room temperature using an inverted microscope (Nikon Eclipse Ts2) with a 20X/0.40 objective (Nikon LWD) equipped with a camera (Nikon DS-Fi3). Each sample was assessed in triplicate and at least 100 cells per well were counted (> 300 cells per experiment).

### RT-qPCR

Cells were treated with 1μM palbociclib for 5 days. RNA was retrotranscribed with High-Capacity cDNA Reverse Transcription Kit (Applied Biosystems, cat#4368814) and 20ng of cDNA amplified using the CFX Connect Real-time PCR system (Bio-Rad) and the PowerUp^TM^ SYBR^TM^ Green Master Mix (Applied Biosystems, cat#A25742) following manufacturer’s instructions. Primers were designed using the Integrated DNA Technologies (IDT) web tool (**Table S8**). Conditions for amplification were: 5 min at 95° C, 40 cycles of 10 sec at 95° C and 7 sec at 62° C. The assay ended with a melting curve program: 15 sec at 95° C, 1 min at 70° C, then ramping to 95° C while continuously monitoring fluorescence. Each sample was assessed in triplicate. Relative quantification was determined to multiple reference genes (human: PSMC4, and B2M and mouse: Rplp0 and Gusb) to ensure reproducibility using the delta-delta CT method.

### *In vivo* mouse experiment

Six-eight week old male athymic NU/J mice were purchased from Jackson Laboratories. All mice were maintained in a HEPA-filtered ventilated rack system at the Animal Facility of the Assembly Building of The Hillman Cancer Center at the University of Pittsburgh School of Medicine. Mice were housed up to 5 mice per cage and in a 12-hour light/dark cycle. All experiments with animals were performed in accordance with institutional guidelines approved by the Institutional Animal Care and Use Committee (IACUC) at the University of Pittsburgh School of Medicine. Ten million SKMEL28 shGFP control or shp16 #1 melanoma cells were subcutaneously injected in the right flank of NU/J mice. Mice were monitored daily to identify palpable tumors, after which mice were randomly assigned to vehicle or methotrexate. Mice were treated by daily intraperitoneal injection of vehicle or 10mg/kg methotrexate (Cayman Chemical Company, cat#13960) diluted in 200μl PBS. Both mice weight and tumors [Length (L) and width (W) (L>W)] were measured 3 times a week. Tumor volume was calculated as ½ (LxW^2^). Animals were euthanized once tumors reached 1000mm^3^. Statistical analysis for tumor burden studies: The data were log-transformed to fit a linear model for tumor growth with fixed effects for mouse type (shp16 vs. control) and for treatment (Vehicle vs MTX). Repeated measures on each subject over time were also specified in the model. Growth rates with standard errors are reported. Growth rate comparisons between groups are presented in the text.

### White blood cell analysis

Fresh whole blood was collected from mice via the inferior vena cava into Microvette K2-EDTA-coated tubes (Sarstedt, cat# 16.444.100). Blood was collected via inferior vena cava using a 20-gauge needle (Becton Dickinson, cat# 305175) after euthanizing mice by co2 inhalation. Total circulating white blood cells (CWBCs) were analyzed using an automated ABAXIS VetScan HM5C Hematology analyzer automated blood counter from (Allied Analytic, Cat# 790-0000).

### Analysis of The Cancer Genome Atlas patient data

Data were extracted from The Cancer Genome Atlas (TCGA) Skin Cutaneous Melanoma Pan-Cancer Atlas (367 metastatic melanomas) using cBioportal survival analysis. Overall survival probability and hazard ratios were calculated using the Log rank statistical test.

### Quantification and Statistical Analysis

GraphPad Prism (version 9.0) and RStudio (Version 2023.06.1+524) were used to perform statistical analysis. Point estimates with standard deviations or standard errors were reported, as indicated, and the appropriate statistical test was performed using all observed experimental data. All statistical tests performed were two-sided and p-values < 0.05 were considered statistically significant.

